# A pan-cancer benchmark of integrated ferroptosis, cuproptosis and disulfidptosis prognostic signatures

**DOI:** 10.64898/2026.06.25.734481

**Authors:** Abdussamed Yasin Demir, Ekrem Yasar

## Abstract

Integrated prognostic signatures combining ferroptosis, cuproptosis, and disulfidptosis are increasingly reported in oncology as advances in risk stratification, yet their added value over simpler pathway-specific or proliferation-related models remains unclear. Here, we developed an integrated regulated cell-death signature and evaluated it through an adversarial pan-cancer benchmark. Using the TCGA pan-cancer cohort comprising 9,808 tumours across 33 cancer types, we curated 118 genes associated with the three cell-death programmes, characterised inter-pathway crosstalk, and derived a 26-gene LASSO-Cox risk signature. The model showed reproducible prognostic performance across cancers, with a pan-cancer concordance index of 0.573 (95% CI, 0.552–0.594), and was independently validated in METABRIC and CGGA cohorts, remaining significant after adjustment for standard clinical variables. However, benchmarking revealed that the integrated signature, although superior to size-matched random gene sets (empirical p < 0.001), did not outperform a ferroptosis-only model (DeLong p = 0.81), indicating no measurable gain from pathway integration. Moreover, much of the prognostic signal reflected tumour proliferation rather than regulated cell death. After adjustment for the proliferation meta-signature (meta-PCNA), ferroptosis performance declined from 0.573 to 0.504, while the integrated model decreased to 0.554. High-risk tumours were more sensitive to anti-proliferative drugs, and the risk score was most strongly associated with E2F, MYC, and G2M target programmes. The signature stratified prognosis but did not predict immune-checkpoint blockade response in IMvigor210 (AUC ≈ 0.50). Importantly, the underlying biology was not merely a modelling artefact. Signature genes showed concordance with protein abundance in CPTAC cohorts, and the three cell-death programmes co-varied within individual malignant cells, with correlations ranging from ρ = 0.46 to 0.66. Overall, our findings indicate that integrated multi-death signatures are reproducible and biologically grounded, yet prognostically redundant and substantially confounded by proliferation. This study provides a cautionary benchmark for the rapidly expanding use of composite regulated cell-death signatures in cancer prognosis.

## 1. Introduction

Regulated cell death (RCD) represents a central determinant of tumour biology, therapeutic vulnerability, and treatment resistance [1–3]. Beyond classical apoptosis, several non-apoptotic cell-death modalities have gained substantial attention in cancer research, particularly those driven by iron, copper, and disulfide stress. Ferroptosis is an iron-dependent form of cell death driven by phospholipid peroxidation and regulated, in part, by the cystine/glutamate antiporter system and the antioxidant enzyme GPX4 [4–7]. Cuproptosis is a copper-dependent cell-death process initiated by the aggregation of lipoylated tricarboxylic-acid-cycle proteins and consequent mitochondrial proteotoxic stress [8–10]. Disulfidptosis, more recently described, results from aberrant disulfide bonding and collapse of the actin cytoskeleton under glucose-starved conditions, particularly in SLC7A11-high cells [11–13]. Notably, shared molecular components, including SLC7A11 and SLC3A2, lie at the interface of these programmes, suggesting that ferroptosis, cuproptosis, and disulfidptosis may not operate as isolated pathways but instead form an interconnected metabolic cell-death axis [14,15].

This mechanistic overlap has encouraged a rapidly growing body of studies that combine genes from two or more of these pathways into integrated prognostic signatures [16–19]. Such models are commonly developed using transcriptomic data from The Cancer Genome Atlas (TCGA) [20,21], often through penalised Cox regression, and are typically presented as improvements in prognostic stratification, tumour-microenvironment characterisation, and prediction of immunotherapy response. However, despite the rapid expansion of these composite signatures across individual cancer types and pan-cancer settings, a key methodological question remains insufficiently addressed: does pathway integration provide measurable value beyond simpler alternatives?

Three controls are particularly important but are often absent. First, an integrated signature should be tested against its constituent single-pathway models. Without this comparison, the apparent prognostic performance of a composite model may simply reflect the contribution of one dominant pathway. Second, a signature of a given size should be evaluated against size-matched random gene sets drawn from cancer-expressed genes, since many transcriptomic signatures may perform well by chance when selected from highly co-regulated cancer-associated genes [22,23]. Third, and most critically, prognostic models derived from bulk tumour transcriptomes are vulnerable to proliferation confounding. Because proliferation is one of the strongest and most pervasive correlates of survival across cancers, an apparent “cell-death” signature may in fact capture generic cell-cycle activity rather than pathway-specific death biology [24].

Here, we address this gap directly. Rather than introducing another integrated cell-death signature as a standalone prognostic tool, we construct a ferroptosis–cuproptosis–disulfidptosis signature within a controlled pan-cancer framework and subject it to an adversarial benchmark. This benchmark includes head-to-head comparisons with single-pathway models, size-matched random-null testing, formal statistical comparison of discriminative performance, and explicit adjustment for proliferation using a meta-PCNA proliferation signature [22]. We further evaluate whether the signature predicts response to immune-checkpoint blockade and assess whether the underlying biology is supported at orthogonal molecular levels, including protein abundance in CPTAC cohorts and pathway co-variation in single-cell malignant populations.

Our findings show that the integrated multi-death signature is reproducible across cohorts and biologically grounded, but its prognostic utility is more limited than its apparent performance suggests. The integrated model does not improve upon the best single-pathway model, is substantially confounded by proliferation, and stratifies prognosis without predicting immunotherapy response. These results provide a cautionary benchmark for the expanding literature on composite regulated cell-death signatures and support the routine inclusion of single-pathway, random-null, and proliferation-adjusted controls in future prognostic signature studies.

## 2. Materials and Methods

### 2.1 Data sources and programmed cell-death gene set

Transcriptomic, clinical, and survival data for the TCGA pan-cancer cohort were obtained from the UCSC Xena platform [25]. The dataset comprised 9,808 tumour samples and 727 adjacent-normal samples across 33 cancer types, using Toil-recomputed RNA-seq expression values represented as log2(TPM + 1) (Table 1; Table ST2). External validation was performed using two independent cohorts: the METABRIC breast cancer cohort (n = 1,979) [26] and the CGGA glioma cohort (n = 657) [27]. For multi-omic validation, proteogenomic datasets from CPTAC were accessed through the cptac Python package [28]. Four CPTAC cohorts were included: clear cell renal cell carcinoma (CCRCC), colon adenocarcinoma (COAD), lung adenocarcinoma (LUAD), and uterine corpus endometrial carcinoma (UCEC). Single-cell RNA-seq validation was conducted using the melanoma dataset GSE72056 [29], which includes pre-annotated malignant, stromal, and immune cell populations.

**Table 1.** Cohort summary. Overview of the transcriptomic data sets used in the study. The TCGA pan-cancer cohort (Toil-recomputed RNA-seq, log2(TPM+1)) spanning 33 cancer types served as the discovery and primary benchmarking set, while METABRIC (breast) and CGGA (glioma) were used as independent external benchmarking cohorts. Numbers of tumour and adjacent-normal samples, cancer types, and tumours with available survival follow-up are listed where applicable. Abbreviations: OS, overall survival; TPM, transcripts per million.

| Cohort | Platform | Tumours<br>(n) | Normals<br>(n) | Cancer<br>types | OS available<br>(n) | Use |
| --- | --- | --- | --- | --- | --- | --- |
| TCGA pan-cancer | RNA-seq (Toil log2 TPM) | 9808 | 727 | 33 | 9777 | discovery + benchmark |
| METABRIC | microarray (Illumina) | — | — | 1 | — | external benchmark |
| CGGA | RNA-seq | — | — | 1 | — | external benchmark |

To evaluate predictive relevance in the immunotherapy setting, two immune-checkpoint blockade cohorts were analysed: IMvigor210, comprising urothelial carcinoma patients treated with anti-PD-L1 therapy [30], and GSE78220, comprising melanoma patients treated with anti-PD-1 therapy [31]. Drug-sensitivity reference data were obtained from GDSC2 [32].

A curated programmed cell-death gene set was assembled for ferroptosis, cuproptosis, and disulfidptosis. The final set contained 118 genes with pathway-membership annotations and literature provenance recorded for each gene (Table ST1). Genes assigned to more than one cell-death programme, such as SLC7A11 and SLC3A2, were explicitly flagged to enable downstream assessment of pathway overlap and crosstalk.

### 2.2 Pathway activity, crosstalk, dysregulation, and molecular subtype analysis

Per-tumour activity of each programmed cell-death programme was quantified using an in-house base-R implementation of single-sample gene-set enrichment analysis (ssGSEA) [33]. ssGSEA scores were calculated separately for ferroptosis, cuproptosis, and disulfidptosis, thereby providing a sample-level estimate of pathway activity across the TCGA pan-cancer cohort. Median pathway activities were also summarised within each cancer type (Table ST3).

Inter-pathway crosstalk was assessed by calculating Spearman correlations between pathway activity scores across the pan-cancer cohort. To complement the transcriptomic co-activity analysis, a protein–protein interaction network for the curated programmed cell-death gene set was retrieved from STRING [34], enabling evaluation of functional connectivity among genes assigned to the three cell-death programmes.

To characterise cancer-associated dysregulation, tumour-versus-normal differential expression analysis was performed across cancer types using matched tumour and adjacent-normal transcriptomic data where available (Supplementary Data SD2). Somatic mutation frequencies of programmed cell-death genes were derived from TCGA mutation calls and summarised at the gene and pathway levels (Table ST4). In addition, aggregate programmed cell-death activity was compared with tumour mutational burden to examine whether global cell-death pathway activity was associated with genomic instability.

Finally, unsupervised consensus clustering was applied to pan-cancer programmed cell-death pathway activity profiles using ConsensusClusterPlus [35]. The resulting molecular subtypes were characterised according to their pathway activity patterns and cancer-lineage composition. Differences in overall survival among subtypes were evaluated using Kaplan–Meier survival analysis and the log-rank test.

### 2.3 Integrated signature, adversarial benchmarking, and external validation

An integrated risk signature was developed using LASSO-penalised Cox regression implemented in glmnet [36] within a pan-cancer training partition. Gene coefficients were estimated by maximising the ℓ1-penalised Cox partial likelihood:

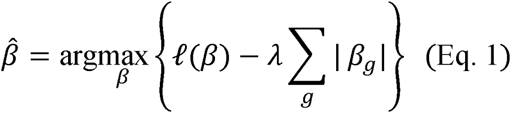

where ℓ(β) denotes the Cox partial log-likelihood, β is the vector of gene coefficients, β_g is the coefficient for gene g, and λ is the regularisation parameter selected by cross-validation (Supplementary Fig. SF5). This procedure yielded a 26-gene model (Table 2). For each sample, a linear risk score was then calculated as:

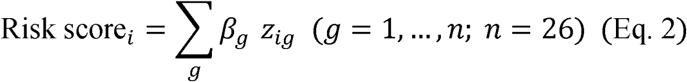

where Risk score_i represents the risk score for sample i, β_g is the LASSO–Cox coefficient of gene g, z_ig is the z-score-standardised expression value of gene g in sample i, and n = 26 denotes the number of genes included in the final signature. The resulting coefficients and risk-score formula were fixed and used unchanged in all downstream analyses. Patients were stratified into high- and low-risk groups using the median risk score as the cut-off. Prognostic discrimination was evaluated using Kaplan–Meier survival analysis with the log-rank test, Harrell’s concordance index (C-index), and time-dependent receiver operating characteristic (ROC) analysis.

**Table 2.**
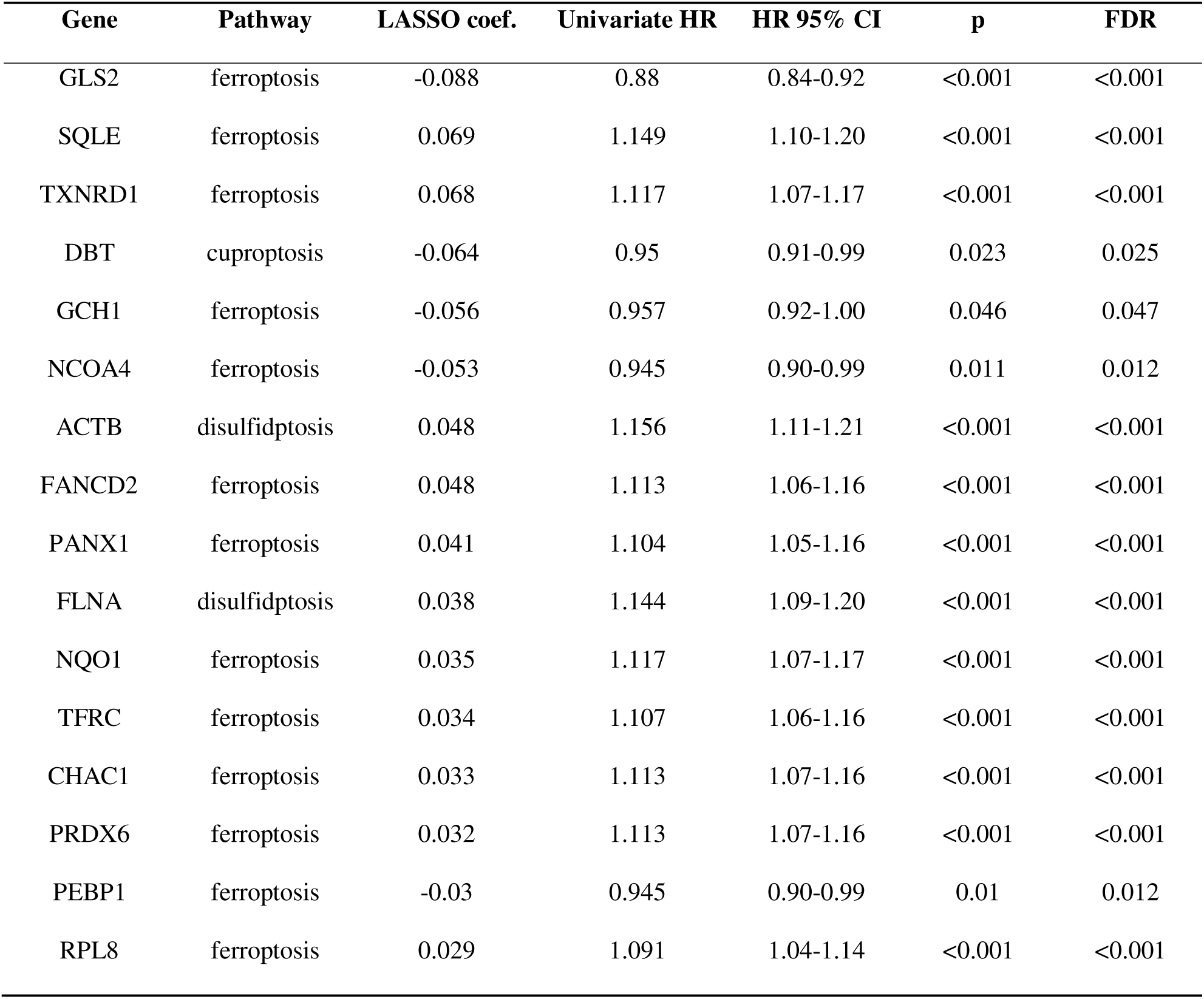
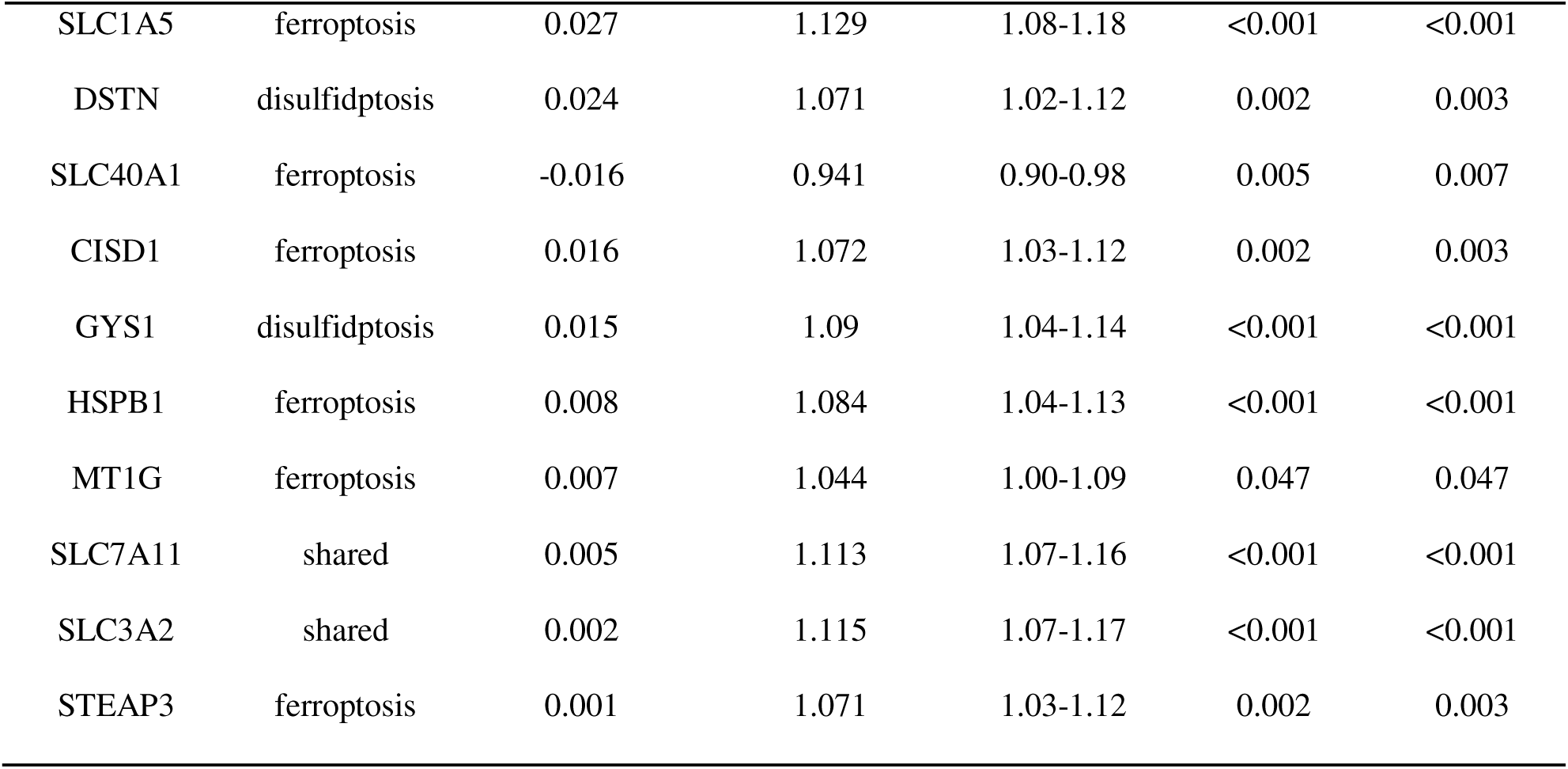
Candidate integrated PCD prognostic signature (26 genes). The 26-gene integrated programmed cell-death signature selected by LASSO-penalised Cox regression in the pan-cancer cohort, jointly drawn from the ferroptosis, cuproptosis and disulfidptosis pathways. For each gene the LASSO coefficient (sign indicates the direction of risk), the pathway of origin, and the univariate pan-cancer Cox hazard ratio with its 95% confidence interval and FDR-adjusted p-value are reported. Genes are ordered by absolute LASSO coefficient. Abbreviations: PCD, programmed cell death; HR, hazard ratio; CI, confidence interval; FDR, false discovery rate.

| Gene | Pathway | LASSO coef. | Univariate HR | HR 95% CI | p | FDR |
| --- | --- | --- | --- | --- | --- | --- |
| GLS2 | ferroptosis | -0.088 | 0.88 | 0.84-0.92 | <0.001 | <0.001 |
| SQLE | ferroptosis | 0.069 | 1.149 | 1.10-1.20 | <0.001 | <0.001 |
| TXNRD1 | ferroptosis | 0.068 | 1.117 | 1.07-1.17 | <0.001 | <0.001 |
| DBT | cuproptosis | -0.064 | 0.95 | 0.91-0.99 | 0.023 | 0.025 |
| GCH1 | ferroptosis | -0.056 | 0.957 | 0.92-1.00 | 0.046 | 0.047 |
| NCOA4 | ferroptosis | -0.053 | 0.945 | 0.90-0.99 | 0.011 | 0.012 |
| ACTB | disulfidptosis | 0.048 | 1.156 | 1.11-1.21 | <0.001 | <0.001 |
| FANCD2 | ferroptosis | 0.048 | 1.113 | 1.06-1.16 | <0.001 | <0.001 |
| PANX1 | ferroptosis | 0.041 | 1.104 | 1.05-1.16 | <0.001 | <0.001 |
| FLNA | disulfidptosis | 0.038 | 1.144 | 1.09-1.20 | <0.001 | <0.001 |
| NQO1 | ferroptosis | 0.035 | 1.117 | 1.07-1.17 | <0.001 | <0.001 |
| TFRC | ferroptosis | 0.034 | 1.107 | 1.06-1.16 | <0.001 | <0.001 |
| CHAC1 | ferroptosis | 0.033 | 1.113 | 1.07-1.16 | <0.001 | <0.001 |
| PRDX6 | ferroptosis | 0.032 | 1.113 | 1.07-1.16 | <0.001 | <0.001 |
| PEBP1 | ferroptosis | -0.03 | 0.945 | 0.90-0.99 | 0.01 | 0.012 |
| RPL8 | ferroptosis | 0.029 | 1.091 | 1.04-1.14 | <0.001 | <0.001 |
| SLC1A5 | ferroptosis | 0.027 | 1.129 | 1.08-1.18 | <0.001 | <0.001 |
| DSTN | disulfidptosis | 0.024 | 1.071 | 1.02-1.12 | 0.002 | 0.003 |
| SLC40A1 | ferroptosis | -0.016 | 0.941 | 0.90-0.98 | 0.005 | 0.007 |
| CISD1 | ferroptosis | 0.016 | 1.072 | 1.03-1.12 | 0.002 | 0.003 |
| GYS1 | disulfidptosis | 0.015 | 1.09 | 1.04-1.14 | <0.001 | <0.001 |
| HSPB1 | ferroptosis | 0.008 | 1.084 | 1.04-1.13 | <0.001 | <0.001 |
| MT1G | ferroptosis | 0.007 | 1.044 | 1.00-1.09 | 0.047 | 0.047 |
| SLC7A11 | shared | 0.005 | 1.113 | 1.07-1.16 | <0.001 | <0.001 |
| SLC3A2 | shared | 0.002 | 1.115 | 1.07-1.17 | <0.001 | <0.001 |
| STEAP3 | ferroptosis | 0.001 | 1.071 | 1.03-1.12 | 0.002 | 0.003 |

To assess whether the integrated signature provided genuine added value, we subjected it to three predefined adversarial controls (Table 3). First, single-pathway signatures were constructed independently from the ferroptosis, cuproptosis, and disulfidptosis gene sets using the same modelling procedure, and their C-indices were compared with that of the integrated model. Differences in discrimination were evaluated using the two-sided DeLong test [37]. Second, a random-null benchmark was generated from 1,000 size-matched random gene sets sampled from cancer-expressed genes. The empirical p-value for the observed integrated-signature C-index was calculated as:

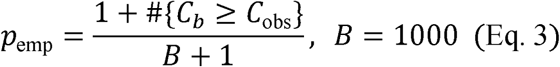

where C_obs is the observed C-index of the integrated signature, C_b is the C-index of the b-th random gene-set model, B = 1,000 is the number of size-matched random gene sets, and #{·} denotes the number of random models satisfying the specified condition. Third, potential proliferation confounding was evaluated by residualising each score on a proliferation meta-signature (meta-PCNA) [22] and re-computing the held-out C-index from the residuals.

**Table 3.**
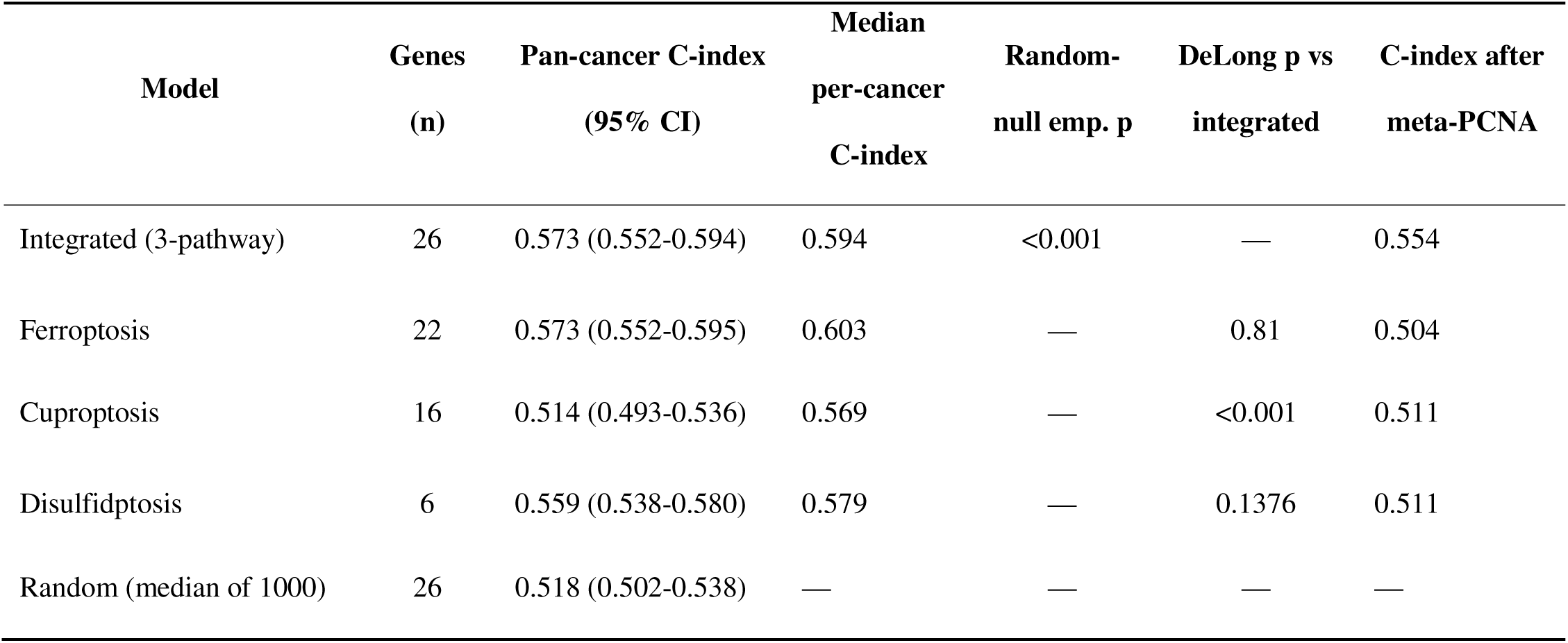
Benchmark of the integrated signature versus single-pathway models, a random-null, and proliferation adjustment. Discrimination of the integrated signature compared with each single-pathway signature, a size-matched random-null, and after adjustment for proliferation. Performance is reported as the pan-cancer Harrell concordance index (with 95% confidence interval) and the median per-cancer concordance index. The integrated signature significantly exceeds the random-null (empirical p < 0.001 against 1000 size-matched random gene sets) but is not superior to the best single pathway (ferroptosis; two-sided DeLong p = 0.81), and its discrimination falls substantially after meta-PCNA proliferation adjustment, indicating that the prognostic signal is largely proliferation-driven. Abbreviations: C-index, Harrell concordance index; CI, confidence interval; emp. p, empirical p-value; meta-PCNA, proliferation meta-signature.

| Model | Genes<br>(n) | Pan-cancer C-index<br>(95% CI) | Median<br>per-cancer<br>C-index | Random-null emp. p | DeLong p vs<br>integrated | C-index after<br>meta-PCNA |
| --- | --- | --- | --- | --- | --- | --- |
| Integrated (3-pathway) | 26 | 0.573 (0.552-0.594) | 0.594 | <0.001 | — | 0.554 |
| Ferroptosis | 22 | 0.573 (0.552-0.595) | 0.603 | — | 0.81 | 0.504 |
| Cuproptosis | 16 | 0.514 (0.493-0.536) | 0.569 | — | <0.001 | 0.511 |
| Disulfidptosis | 6 | 0.559 (0.538-0.580) | 0.579 | — | 0.1376 | 0.511 |
| Random (median of 1000) | 26 | 0.518 (0.502-0.538) | — | — | — | — |

For external validation, the frozen 26-gene signature, including its original coefficients, was applied without refitting to the METABRIC and CGGA cohorts. Prognostic performance was assessed using Kaplan–Meier/log-rank analysis, univariate Cox regression, and clinically adjusted Cox models. Adjustment covariates were age, grade, and stage for METABRIC, and age, grade, IDH status, and sex for CGGA. Cohort-specific hazard ratios were subsequently combined using fixed-effect meta-analysis (Table ST6).

### 2.4 Tumour microenvironment, drug sensitivity, and functional enrichment analyses

Tumour microenvironment composition was assessed using the ESTIMATE algorithm [38], which was used to infer stromal and immune content from bulk transcriptomic profiles. Immune-cell infiltration was further quantified using marker-based immune signatures described by Danaher et al. [39]. Associations between the risk score and stromal/immune scores, immune-cell infiltration estimates, functional immune signatures, and immune-checkpoint gene expression were evaluated using Spearman correlation. Multiple testing was controlled using the Benjamini–Hochberg false discovery rate (FDR) procedure (Table ST7).

To determine whether the signature was predictive of immune-checkpoint blockade response, two independent immunotherapy cohorts were analysed. In each cohort, the ability of the risk score to discriminate responders from non-responders was quantified using receiver operating characteristic analysis and the area under the curve (ROC AUC). Associations between the risk score and on-treatment survival were evaluated using Cox proportional hazards regression (Table ST8). Drug-sensitivity associations were assessed using reference pharmacogenomic data from GDSC2. Drug response was imputed by ridge regression implemented in glmnet [36], and predicted drug sensitivity was compared between high- and low-risk groups. Differences in predicted natural-log-transformed IC50 values were tested using the Wilcoxon rank-sum test, with FDR correction applied across drugs (Table ST11).

Finally, functional enrichment analysis was performed to identify biological programmes associated with the risk score. Genes were ranked according to their correlation with the risk score, and pre-ranked gene-set enrichment analysis was conducted using fgsea [40]. Enrichment was tested against the Hallmark and Gene Ontology Biological Process collections from MSigDB [41,42]. Full enrichment results are provided in Supplementary Data SD3.

### 2.5 Multi-omic and single-cell validation

To evaluate whether the transcriptome-derived signature was supported at the protein level, multi-omic validation was performed using CPTAC cohorts. For each signature gene, mRNA–protein concordance was quantified using Spearman correlation between transcript abundance and matched protein abundance. In addition, RNA-based and protein-based risk scores were calculated and compared to assess whether the risk signal was preserved across molecular layers (Table ST9). Single-cell validation was performed using the GSE72056 dataset. Pathway activity was quantified at the single-cell level by calculating module scores for each regulated cell-death programme without prior clustering. For each gene, log-normalised expression values were standardised across cells, and the module score for each cell was defined as the mean standardised expression of genes belonging to the corresponding pathway:

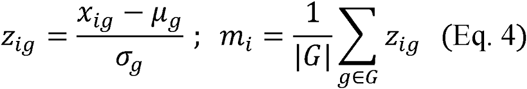

where x_ig denotes the log-normalised expression of gene g in cell i, μ_g and σ_g represent the mean and standard deviation of gene g across all cells, z_ig is the resulting standardised expression value, m_i is the pathway module score for cell i, and G denotes the set of genes assigned to the pathway.

These module scores were used to characterise cell-type-specific pathway activity and to quantify pathway co-variation within malignant cells using Spearman correlation. Summary results are provided in Table ST10, and per-tumour single-cell module scores are reported in Supplementary Data SD1.

### 2.6 Statistical analysis and reproducibility

All statistical analyses were performed in R 4.6.0 with Bioconductor 3.23. Python was used for CPTAC data access and preprocessing where required. Unless otherwise stated, statistical tests were two-sided, and multiple comparisons were adjusted using the Benjamini–Hochberg false discovery rate procedure. Statistical significance was defined as an adjusted or nominal p < 0.05, as appropriate for each analysis.

To ensure reproducibility, all analysis code and processed data have been deposited in public repositories. The code required to reproduce the analyses is available through GitHub, while processed result tables, intermediate analysis checkpoints, and large supplementary data files, including Supplementary Data SD1–SD3, have been archived on Zenodo. Together, these resources provide the computational workflow, processed datasets, and outputs necessary to reproduce the analyses reported in this study. Further details are provided in the Data and Code Availability section.

## 3. Results

### 3.1 A coordinated ferroptosis–cuproptosis–disulfidptosis crosstalk axis across cancers

A curated set of 118 programmed cell-death genes was assembled to represent ferroptosis, cuproptosis, and disulfidptosis, with pathway-membership annotations and literature provenance recorded for each gene (Table ST1). Among these genes, SLC7A11 and SLC3A2 were shared across pathways, consistent with their proposed role at the metabolic interface of these cell-death programmes (Fig 1A). Pathway activity was quantified across the TCGA pan-cancer cohort using single-sample gene-set enrichment analysis (ssGSEA). This cohort included 9,808 tumours across 33 cancer types (Table 1; per-cancer sample counts in Table ST2; full sample overview in Supplementary Fig SF1). The three pathway activity scores were positively and significantly correlated across tumours (Fig 1B), indicating that ferroptosis, cuproptosis, and disulfidptosis tend to co-vary rather than behave as independent programmes.

**Figure 1.**
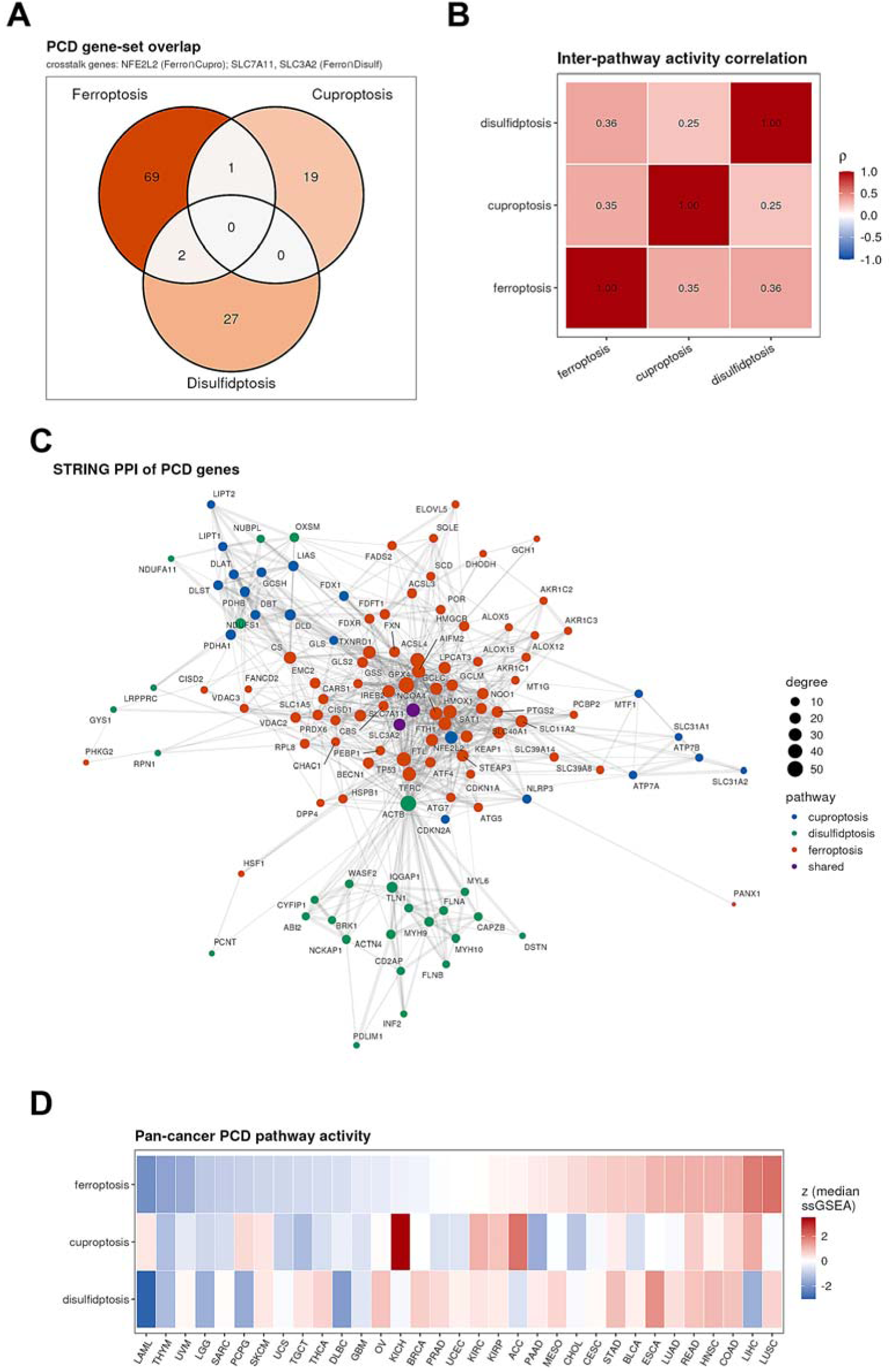
A coordinated ferroptosis–cuproptosis–disulfidptosis crosstalk axis across cancers. (A) Overlap of the curated 118-gene programmed cell-death (PCD) set across the three pathways. (B) Pairwise Spearman correlation of per-tumour ssGSEA pathway-activity scores across the TCGA pan-cancer cohort. (C) STRING protein–protein interaction network of the PCD gene set. (D) Distribution of PCD pathway activity across tumour lineages.

The structural organisation of the curated gene set was further examined using a STRING protein–protein interaction network. The resulting network was densely connected, with shared transporter-related genes, including SLC7A11 and SLC3A2, positioned among the major hubs (Fig 1C). Despite this coordinated architecture, pathway activity varied substantially across tumour lineages (Fig 1D; per-cancer median activities in Table ST3), suggesting cancer-type-specific engagement of the cell-death axis. Somatic mutation frequencies of the constituent genes were generally low at the pan-cancer level (Table ST4), indicating that variation in pathway activity is more likely to reflect transcriptional or regulatory differences than recurrent genetic alteration. Together, these findings support the presence of a coordinated ferroptosis–cuproptosis–disulfidptosis axis across human cancers and provide a biologically grounded basis for subsequent prognostic and benchmarking analyses.

### 3.2 Pan-cancer dysregulation and molecular subtypes of the PCD axis

The programmed cell-death (PCD) axis showed broad dysregulation across human cancers. Tumour-versus-normal differential expression analysis revealed widespread up- and down-regulation of PCD genes across cancer types (Fig 2A; per-gene log2 fold-change across cancers in Supplementary Fig SF2; full results in Supplementary Data SD2). The extent of dysregulation varied by tumour lineage, as reflected by differences in the number of differentially expressed genes across cancer types (Fig 2B). Somatic alterations in PCD genes were recurrent but generally modest at the pan-cancer level, as shown by the oncoplot analysis (Fig 2C). In addition, aggregate PCD activity was associated with tumour mutational burden, suggesting a relationship between cell-death pathway activity and broader genomic instability (Fig 2D).

**Figure 2.**
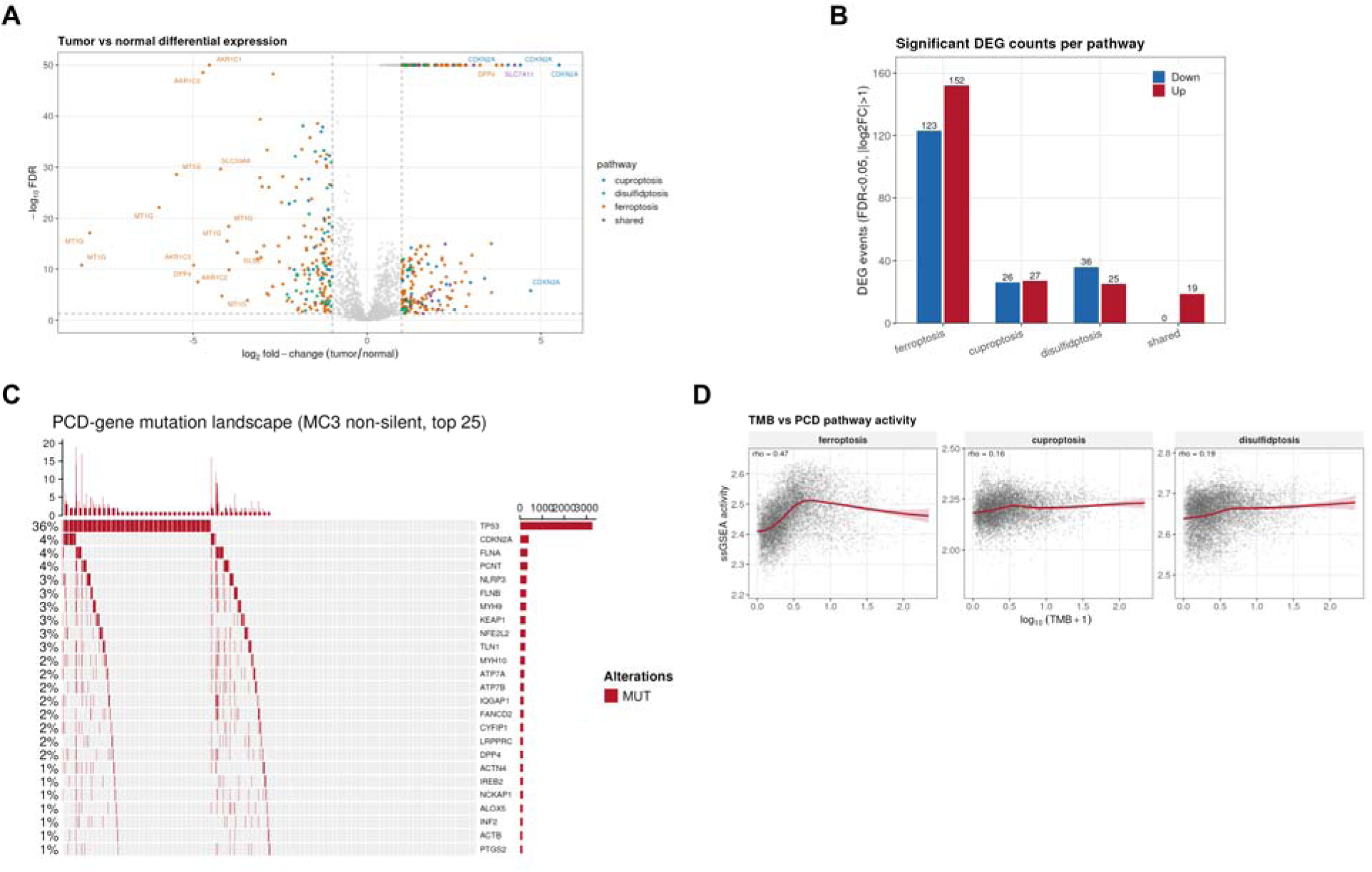
Pan-cancer dysregulation of the PCD axis. (A) Volcano plot of tumour-versus-normal differential expression of PCD genes. (B) Number of differentially expressed PCD genes per cancer type. (C) Oncoplot of recurrent somatic mutations in PCD genes. (D) Relationship between aggregate PCD pathway activity and tumour mutational burden.

Unsupervised consensus clustering based on pan-cancer PCD activity identified robust molecular subtypes (Fig 3A,B). These subtypes differed in pathway activity profiles (Fig 3C), global gene-expression patterns (Supplementary Fig SF4), and tumour-lineage composition (Fig 3D), indicating that the PCD axis captures biologically meaningful heterogeneity across cancers. However, the identified PCD subtypes did not significantly stratify overall survival (Supplementary Fig SF3). This negative result is important, as it indicates that unsupervised structure within the PCD axis is not, by itself, strongly prognostic. This observation provided the rationale for subsequent supervised modelling and adversarial benchmarking of a prognostic cell-death signature.

**Figure 3.**
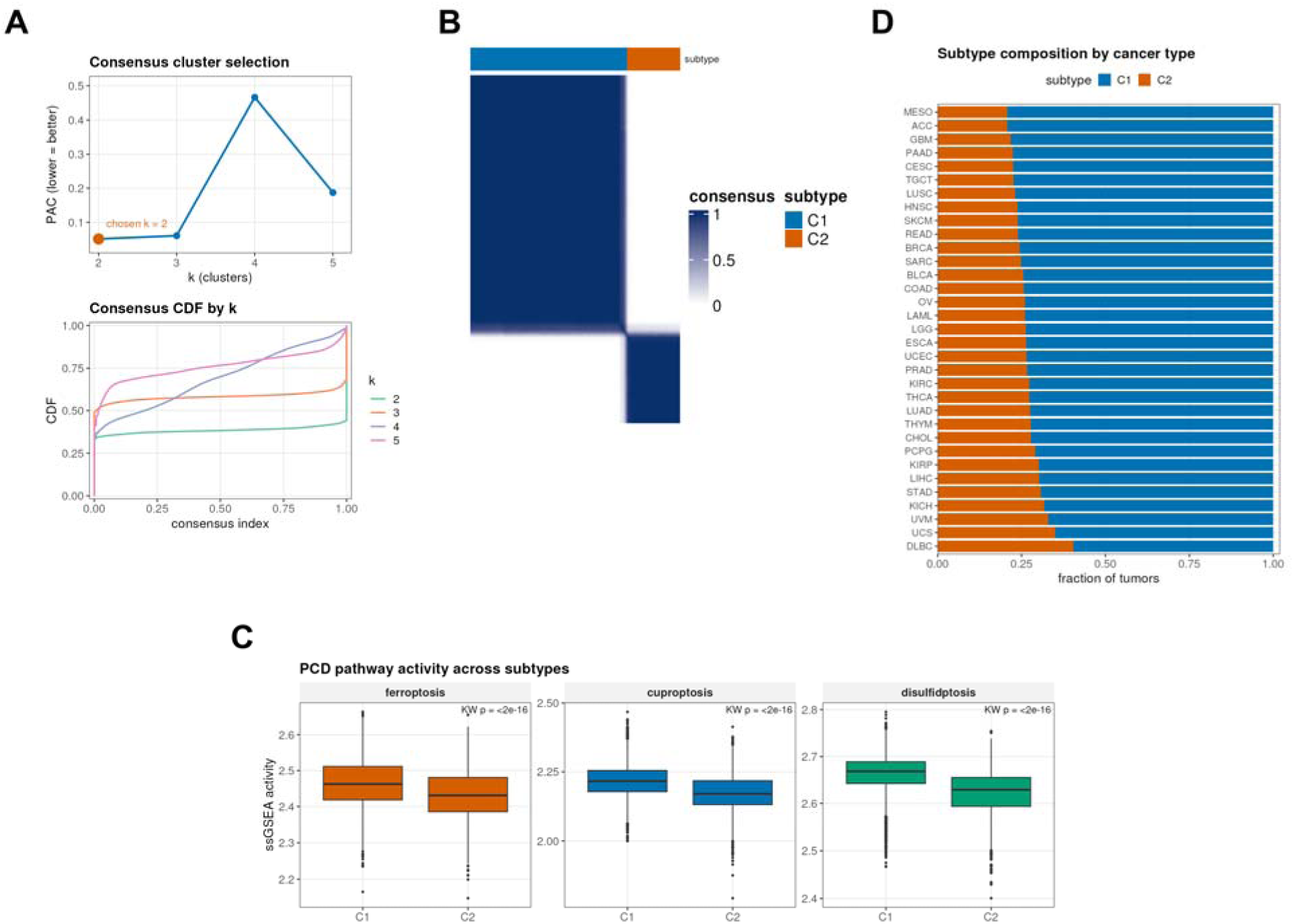
Programmed cell-death molecular subtypes. (A) Consensus clustering diagnostics used to select the number of subtypes. (B) Consensus matrix at the selected number of clusters. (C) PCD pathway activity across subtypes. (D) Cancer-lineage composition of the subtypes.

### 3.3 An integrated 26-gene prognostic signature

To generate an integrated prognostic model under a best-case scenario, a LASSO-penalised Cox regression model was trained using the pan-cancer cohort. This analysis yielded a frozen 26-gene risk signature comprising genes from ferroptosis, cuproptosis, and disulfidptosis, together with shared transporter components located at the interface of these pathways (Table 2; cross-validation shown in Supplementary Fig SF5).

The resulting risk score stratified patients into distinct survival groups in both the training and held-out test partitions (Supplementary Fig SF6, SF7). Time-dependent ROC analysis further showed discriminatory performance above chance across evaluated time points (Supplementary Fig SF8). In the held-out test data, the signature achieved a pan-cancer concordance index of 0.573 (95% CI, 0.552–0.594), with a median per-cancer C-index of 0.594 (Fig 4A; per-cancer estimates in Table ST5). According to the conventional criteria used in many prognostic-signature studies, these results would support the model as a successful pan-cancer risk signature. This made the 26-gene model an appropriate and informative candidate for subsequent adversarial benchmarking against single-pathway, random-null, and proliferation-adjusted controls.

**Figure 4.**
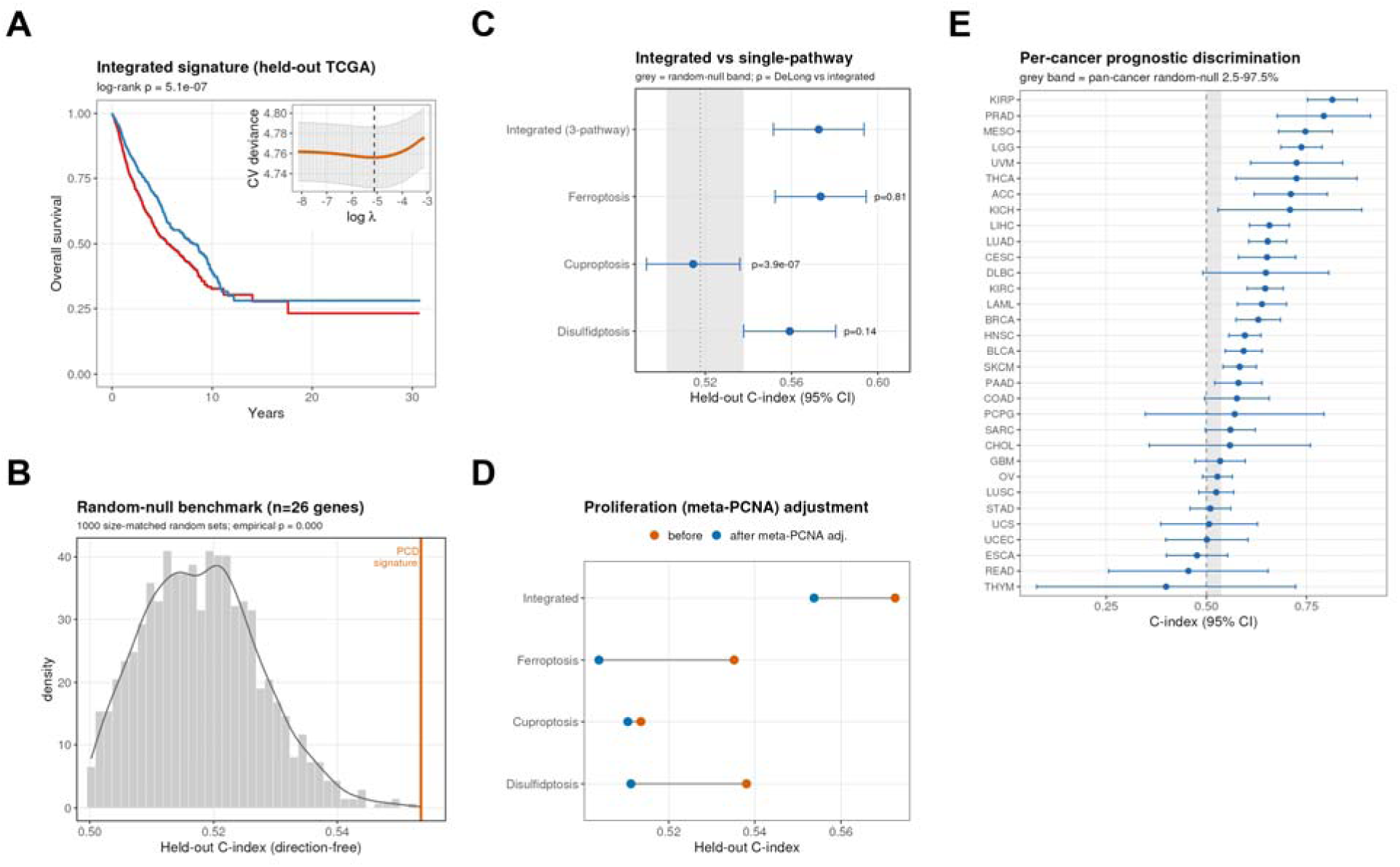
Adversarial benchmark of the integrated PCD signature. (A) Per-cancer concordance index (C-index) of the integrated 26-gene signature. (B) Observed integrated C-index relative to the size-matched random-null distribution. (C) Head-to-head C-index comparison of the integrated signature with single-pathway models (two-sided DeLong test). (D) C-index before and after meta-PCNA proliferation adjustment. (E) Forest plot of per-cancer C-index estimates.

### 3.4 Benchmark analysis shows that the integrated signature adds no value over the strongest single pathway

The integrated 26-gene signature was next evaluated against three predefined adversarial controls (Fig 4; Table 3). In comparison with 1,000 size-matched random gene sets, the integrated signature performed significantly above the random-null distribution (empirical p < 0.001; Fig 4B). This result indicates that the model captures a genuine cancer-relevant prognostic signal rather than random noise.

However, head-to-head comparison with single-pathway models showed that pathway integration did not improve prognostic discrimination. A ferroptosis-only signature achieved the same pan-cancer C-index as the integrated model (0.573), and the difference between the two models was not statistically significant (two-sided DeLong p = 0.81; Fig 4C). Cuproptosis and disulfidptosis models showed weaker individual performance, with C-indices of 0.514 and 0.559, respectively. Nevertheless, the strongest constituent pathway matched the performance of the full integrated signature. Per-cancer C-index estimates further supported this conclusion. The forest plot of cancer-specific performance showed generally consistent but modest discrimination across tumour lineages, without identifying any cancer type in which the integrated model was decisively superior to the single-pathway alternatives (Fig 4E). Thus, although the integrated signature clearly exceeded random expectation, combining ferroptosis, cuproptosis, and disulfidptosis did not provide measurable prognostic gain beyond the strongest individual pathway. This finding indicates that the apparent success of the integrated model is largely redundant with ferroptosis-associated signal.

### 3.5 External multi-cohort validation

Although the integrated signature did not provide incremental prognostic value over the strongest single-pathway model, its prognostic signal was reproducible across independent external cohorts. The frozen 26-gene risk score was applied without refitting to METABRIC, a breast cancer cohort comprising 1,979 patients, and CGGA, a glioma cohort comprising 657 patients. In both cohorts, the risk score significantly stratified patients into distinct survival groups by Kaplan–Meier analysis and log-rank testing (Fig 5A). Cox regression analyses further confirmed the prognostic relevance of the signature. The risk score remained significantly associated with survival after adjustment for standard clinical covariates, with adjusted hazard ratios of 1.12 in METABRIC (p = 0.002) and 1.19 in CGGA (p = 0.002). In fixed-effect meta-analysis of cohort-specific univariate estimates, the pooled hazard ratio was 1.22 (95% CI, 1.16–1.29), indicating a consistent adverse association between higher risk score and survival outcome across validation settings (Fig 5B; Table ST6).

**Figure 5.**
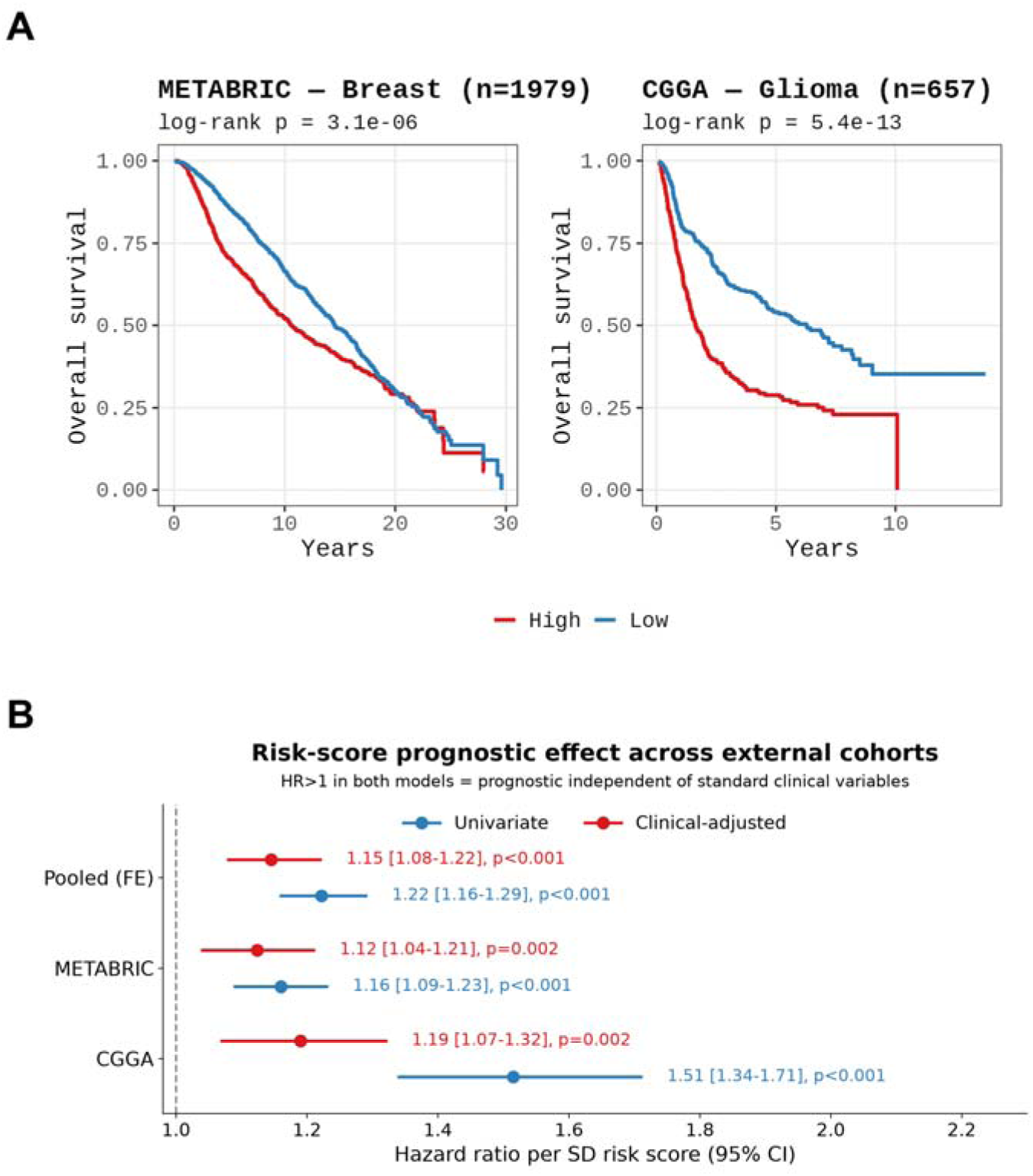
External multi-cohort validation. (A) Kaplan–Meier overall survival of risk groups defined by the frozen signature in the METABRIC (breast) and CGGA (glioma) cohorts. (B) Forest plot of univariate and clinically adjusted hazard ratios per cohort, with the fixed-effect pooled estimate.

These findings demonstrate that the integrated signature captures a stable and reproducible prognostic signal across distinct tumour lineages and data platforms. However, in light of the adversarial benchmark analysis, reproducibility alone should not be interpreted as evidence of added clinical utility. Rather, the external validation supports the robustness of the signal while reinforcing the distinction between prognostic reproducibility and incremental value beyond simpler pathway-specific models.

### 3.6 Tumor microenvironment and immunotherapy: prognostic but not predictive

The integrated risk score was associated with selected features of tumour microenvironment biology. Higher risk scores correlated with ESTIMATE-derived stromal scores and the overall ESTIMATE score, whereas the association with the immune score was weaker and did not remain significant after multiple-testing correction (Fig 6A). Marker-based immune-infiltration analysis further showed positive correlations with dendritic cells, macrophages, neutrophils, and regulatory T cells, while negative correlations were observed with mast cells and CD8 T cells (Fig 6B). The risk score was also associated with functional immune signatures and immune-checkpoint gene expression, indicating that the signature captures variation in immune-related tumour states despite not being uniformly associated with global immune content (Fig 6C,D; Table ST7). However, these immune-associated features did not translate into prediction of immune-checkpoint blockade benefit. In the IMvigor210 cohort of anti-PD-L1-treated urothelial carcinoma and the GSE78220 cohort of anti-PD-1-treated melanoma, the risk score failed to discriminate responders from non-responders. In IMvigor210, response classification was essentially uninformative, with an AUC of approximately 0.50. Consistently, the risk score did not significantly stratify on-treatment survival in IMvigor210, the cohort with available survival data (Fig 6E,F; Table ST8).

**Figure 6.**
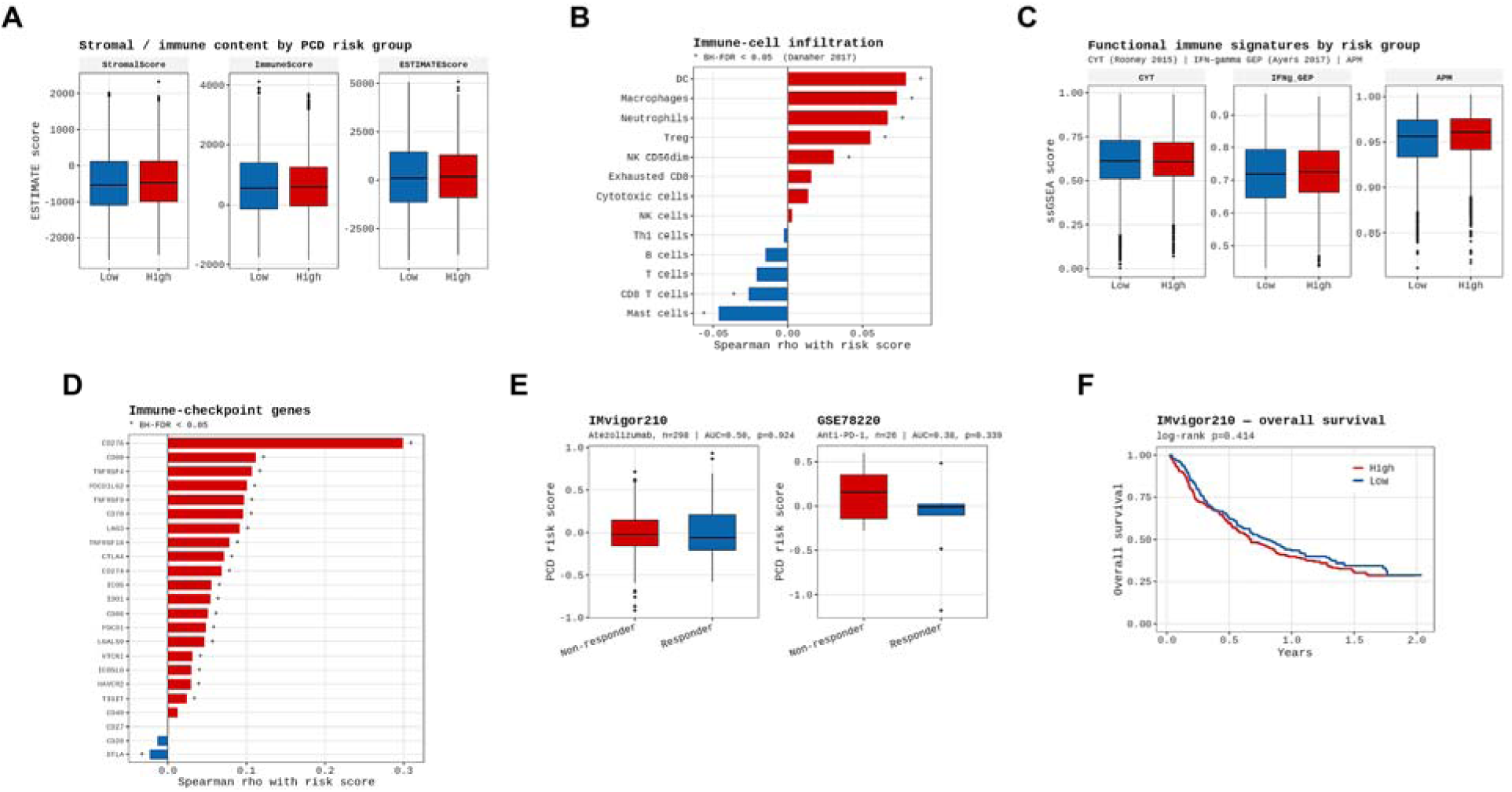
Tumour microenvironment and immunotherapy. (A) Correlation of the risk score with ESTIMATE stromal and immune scores. (B) Association of the risk score with immune-cell infiltration (Danaher signatures). (C) Functional immune signatures. (D) Immune-checkpoint gene expression. (E) Discrimination of immune-checkpoint-blockade (ICB) responders (ROC) and (F) on-treatment survival in the IMvigor210 and GSE78220 cohorts.

These findings indicate that the integrated signature is associated with aspects of tumour microenvironment organisation and immune-related transcriptional states, but does not function as a predictive biomarker of immune-checkpoint blockade response. Thus, the signature should be interpreted as prognostic rather than predictive, underscoring an important distinction that is often insufficiently separated in transcriptomic signature studies.

### 3.7 Crosstalk biology is supported at protein and single-cell resolution

Although the integrated signature did not provide incremental prognostic value over the strongest single-pathway model, orthogonal validation supported the biological relevance of the underlying crosstalk axis. In CPTAC proteogenomic cohorts, signature genes showed concordance between mRNA and protein abundance, with median per-gene Spearman correlations ranging from ρ = 0.38 to 0.62 across CCRCC, COAD, LUAD, and UCEC. Among these genes, SLC7A11 showed particularly strong mRNA–protein concordance, with a correlation of approximately ρ = 0.85. RNA-based and protein-based risk scores were also positively correlated within each cohort, with Spearman correlations ranging from ρ = 0.43 to 0.61 (Fig 7A,B; Table ST9).

**Figure 7.**
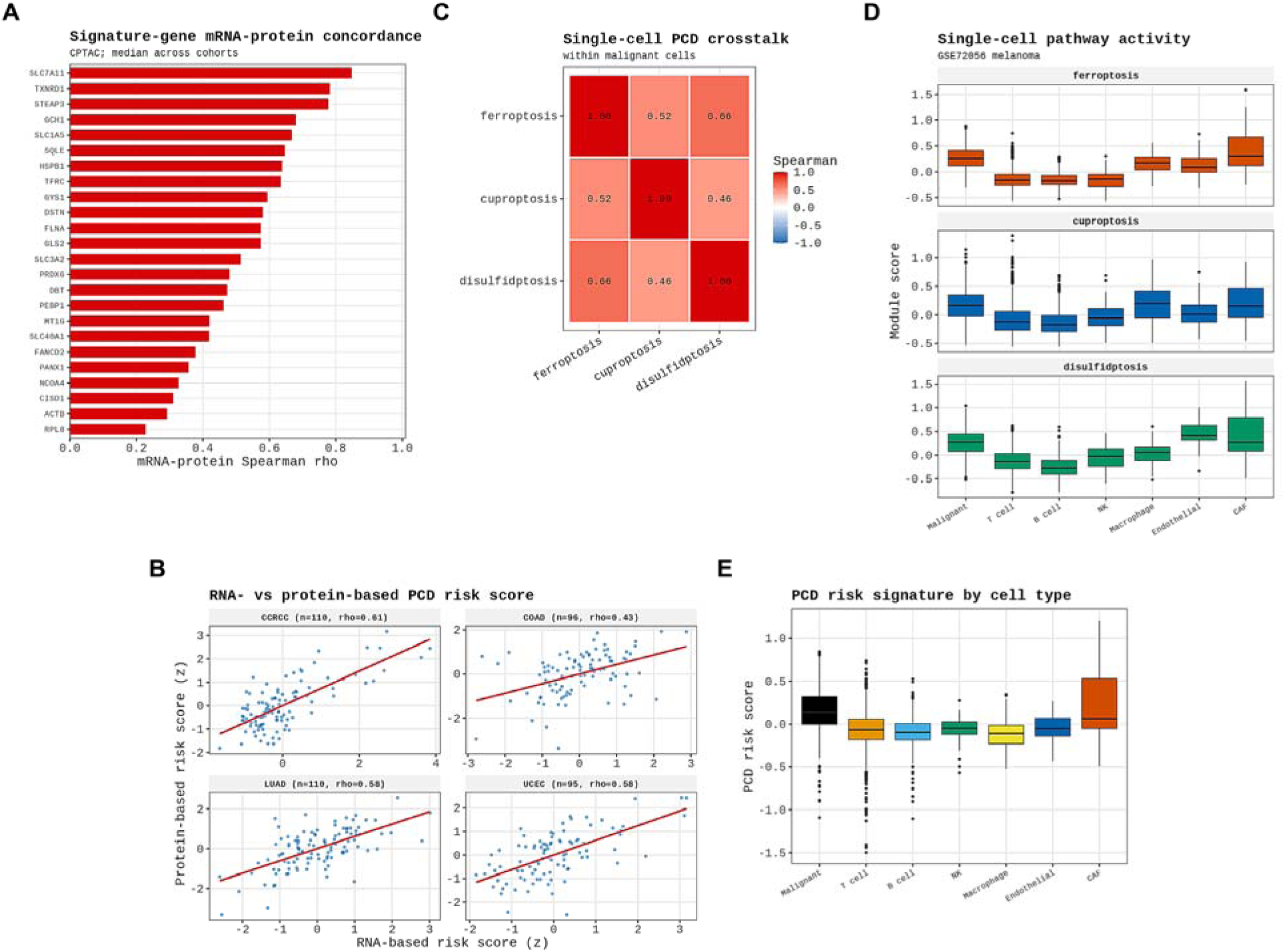
Multi-omic and single-cell validation. (A) Per-gene mRNA–protein concordance of signature genes across CPTAC cohorts. (B) Agreement between RNA-based and protein-based risk scores. (C–E) Single-cell analysis of melanoma (GSE72056): pathway activity by cell type, within-malignant-cell pathway crosstalk, and risk score across cell types.

Single-cell analysis further supported the presence of coordinated pathway activity at cellular resolution. In the melanoma single-cell RNA-seq dataset GSE72056, ferroptosis, cuproptosis, and disulfidptosis module scores co-varied within individual malignant cells, with Spearman correlations ranging from ρ = 0.46 to 0.66. This finding recapitulates the bulk-tissue crosstalk pattern at the level of individual tumour cells. Malignant cells and cancer-associated fibroblasts showed the highest pathway activities and risk scores, indicating that the PCD axis is not uniformly distributed across the tumour microenvironment but is enriched in specific cellular compartments (Fig 7C–E; Table ST10; per-tumour pathway scores in Supplementary Data SD1). Together, these multi-omic and single-cell analyses indicate that the ferroptosis–cuproptosis–disulfidptosis crosstalk axis represents a genuine, multi-scale biological phenomenon. This biological validity makes the observed prognostic redundancy particularly important, demonstrating that a signature can capture real tumour biology without necessarily providing added prognostic value beyond simpler pathway-specific models.

### 3.8 The prognostic signal is largely proliferation-driven

Three independent lines of evidence indicated that the residual prognostic signal of the integrated signature was driven predominantly by proliferation rather than cell-death-specific biology. First, adjustment for the proliferation meta-signature meta-PCNA substantially reduced model discrimination. The C-index of the integrated model decreased from 0.573 to 0.554, while the ferroptosis-only model declined more markedly, from 0.573 to 0.504 (Fig 4D; Table 3). This reduction suggests that a substantial component of the apparent prognostic performance overlaps with proliferation-associated transcriptional activity. Second, functional enrichment analysis of genes ranked by correlation with the risk score identified proliferation-related programmes as the dominant biological signal. Among Hallmark gene sets, the strongest enrichments included E2F_TARGETS, MYC_TARGETS, G2M_CHECKPOINT, and MTORC1_SIGNALING (Fig 8C). Gene Ontology Biological Process enrichment showed a similar pattern, with top terms dominated by rRNA processing, DNA replication, chromosome segregation, and related cell-cycle-associated processes (Fig 8D; full enrichment results in Supplementary Data SD3). These transcriptional patterns indicate that higher risk scores are closely aligned with proliferative tumour states. Third, predicted drug-sensitivity analysis using GDSC2 showed that high-risk tumours were preferentially sensitive to anti-proliferative cytotoxic agents, including docetaxel, paclitaxel, vinblastine, and AZD5991 (Fig 8A,B; Table ST11). This pharmacological profile further supports the interpretation that the risk score captures proliferative vulnerability rather than a pathway-specific regulated cell-death dependency.

**Figure 8.**
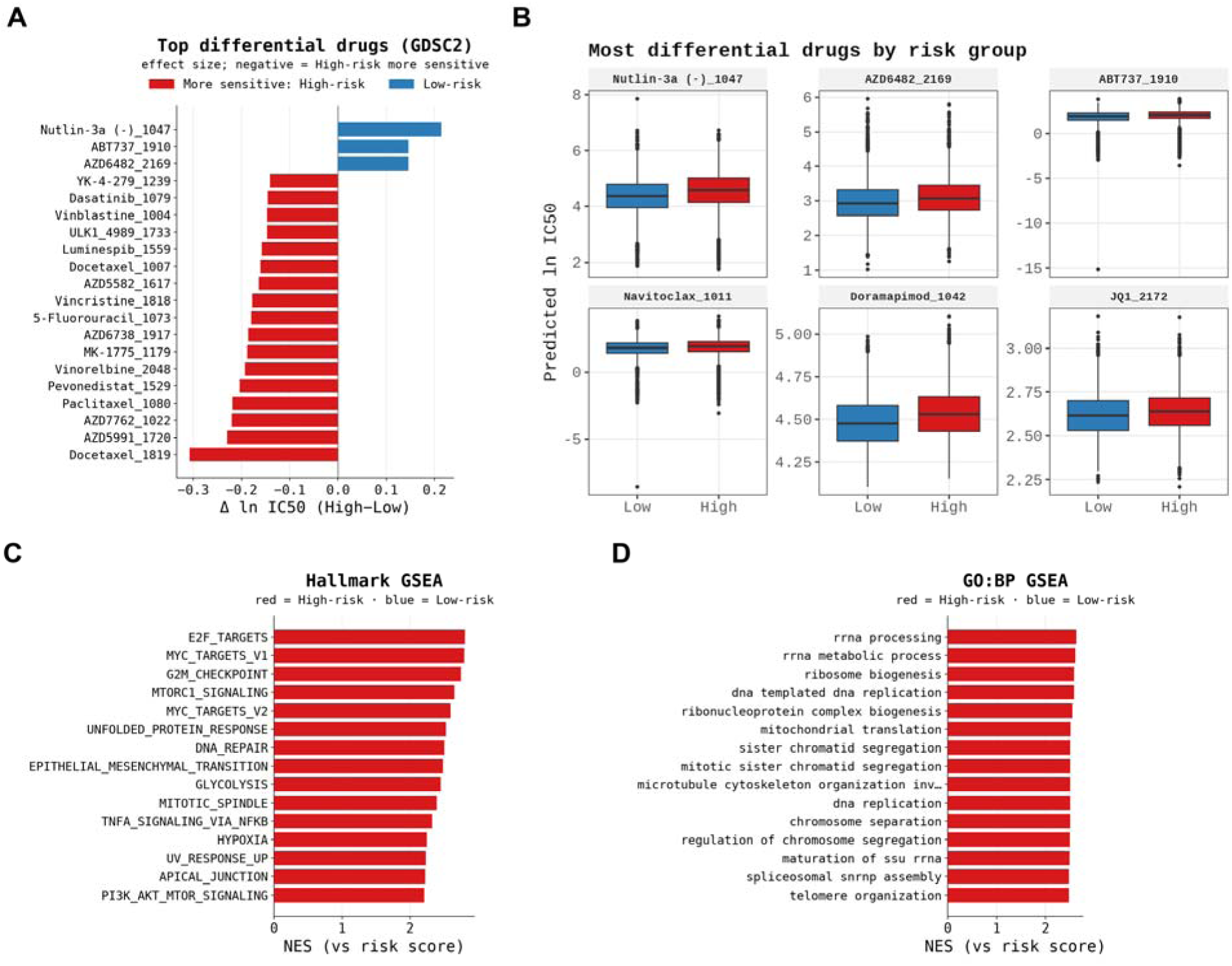
Drug sensitivity and functional enrichment of the risk score. (A) Compounds most differentially sensitive between risk groups. (B) Predicted IC50 distributions for representative anti-proliferative drugs. (C) Hallmark gene-set enrichment of the risk score. (D) GO Biological Process enrichment.

Together, these statistical, transcriptional, and pharmacological findings converge on the same conclusion: although the signature is reproducibly prognostic, a substantial part of its prognostic signal behaves as a proliferation proxy rather than a specific readout of ferroptosis–cuproptosis–disulfidptosis biology.

## 4. Discussion

This study evaluated whether integrating ferroptosis, cuproptosis, and disulfidptosis into a single prognostic signature provides added value across human cancers when tested against appropriate controls. The answer is nuanced but clear. The integrated 26-gene signature was reproducible across independent cohorts and was supported by multi-scale biological evidence, yet it did not improve prognostic discrimination beyond the strongest constituent pathway. Moreover, a substantial component of its prognostic signal reflected tumour proliferation rather than regulated cell-death-specific biology. These findings indicate that reproducibility and biological plausibility, although necessary, are not sufficient to establish incremental clinical value. Without direct comparison to single-pathway models, random-null benchmarking, and explicit proliferation adjustment, integrated prognostic signatures may appear robust while remaining redundant or confounded [22,23].

The biological basis for a ferroptosis–cuproptosis–disulfidptosis crosstalk axis was strongly supported. The curated gene set revealed shared pathway components, particularly the transporter-related genes SLC7A11 and SLC3A2, which lie at the metabolic interface of these cell-death programmes [12,14]. Across the TCGA pan-cancer cohort, pathway activities were positively correlated, and the corresponding protein–protein interaction network showed dense functional connectivity. This coordinated structure was also observed beyond bulk transcriptomics. In CPTAC proteogenomic cohorts, signature genes showed concordance between mRNA and protein abundance, with SLC7A11 among the most concordant genes. At single-cell resolution, ferroptosis, cuproptosis, and disulfidptosis module scores co-varied within individual malignant melanoma cells, recapitulating the bulk-tissue crosstalk pattern at cellular resolution. These findings indicate that the integrated axis is not merely a statistical artefact but represents a genuine, conserved feature of tumour biology. This is consistent with the established role of the system x_c_^−^/SLC7A11 axis in tumour metabolism and immune interaction, including the ability of CD8^+^ T-cell-derived interferon-γ to suppress cystine uptake and promote ferroptosis [43].

However, the adversarial benchmark substantially reframed the prognostic interpretation. Integrated cell-death signatures are frequently presented as advances in risk stratification [16–19], but the present analysis showed that integration itself did not provide measurable benefit. Although the integrated signature significantly outperformed size-matched random gene sets, a ferroptosis-only model achieved the same pan-cancer C-index, with no significant difference by DeLong testing. This suggests that the prognostic information captured by the integrated model is largely contained within the strongest individual pathway, rather than emerging from the combination of ferroptosis, cuproptosis, and disulfidptosis. The shared transporter machinery and coordinated pathway activity may therefore produce overlapping prognostic signals, limiting the incremental value of composite pathway integration.

A second major finding was that the residual prognostic signal was substantially proliferation-driven. Adjustment for the meta-PCNA proliferation signature reduced the performance of the integrated model and drove single-pathway models, particularly ferroptosis, close to random discrimination. Functional enrichment analysis further showed that genes correlated with the risk score were dominated by proliferation-associated programmes, including E2F targets, MYC targets, G2M checkpoint, mTORC1 signalling, DNA replication, chromosome segregation, and ribosome biogenesis. The pharmacological profile was consistent with this interpretation: high-risk tumours were preferentially sensitive to anti-proliferative cytotoxic agents such as docetaxel, paclitaxel, and vinblastine. Together, these statistical, transcriptional, and pharmacological findings indicate that the signature behaves substantially as a proliferation proxy. This is particularly important because proliferation is among the strongest generic correlates of survival in bulk tumour transcriptomes [22,24], and many highly expressed cancer-associated genes are coupled to cell-cycle activity. As a result, a transcriptomic “cell-death” risk score may capture proliferative tumour state more strongly than pathway-specific regulated cell-death biology. These observations align with broader concerns regarding the reproducibility, redundancy, and over-interpretation of transcriptomic prognostic signatures [23,44].

The distinction between prognostic and predictive value is also critical. The integrated risk score was associated with tumour microenvironment organisation, stromal content, immune-cell infiltration patterns, functional immune signatures, and immune-checkpoint gene expression. Such associations are often used to motivate immunotherapy relevance in signature studies [45,46]. However, immune association did not translate into prediction of immune-checkpoint blockade benefit. The score failed to discriminate responders from non-responders in IMvigor210 and GSE78220, with response classification in IMvigor210 close to random performance. In addition, the risk score did not significantly stratify on-treatment survival in IMvigor210, the cohort with available survival data. These findings support the interpretation that the signature is prognostic but not predictive. Prediction of immunotherapy benefit remains more appropriately evaluated using response-linked biomarkers, such as tumour mutational burden, interferon-γ-related T-cell-inflamed signatures, antigen presentation markers, and other treatment-specific features [47–49]. Prognostic association with immune features should therefore not be assumed to imply predictive utility.

These results have direct methodological implications for the rapidly expanding literature on integrated regulated cell-death signatures. Many such studies report composite signatures as positive prognostic advances, but the controls applied here are rarely included. Three benchmarks should become routine. First, integrated signatures should be compared directly against the best constituent single-pathway model to establish whether integration adds discriminatory value. Second, performance should be tested against size-matched random gene sets drawn from an appropriate background of cancer-expressed genes. Third, proliferation adjustment should be included explicitly, given the strong and pervasive influence of cell-cycle activity on survival-associated transcriptomic models. These controls require no additional experimental data, are computationally straightforward, and would help prevent the accumulation of reproducible but redundant prognostic models; they also complement established reporting standards for prognostic-model and tumour-marker studies [50,51]. In this context, the present study provides a reusable benchmark framework rather than simply another signature model.

Several strengths support the interpretation of these findings. The analysis was performed in a large TCGA pan-cancer cohort, externally validated using frozen coefficients in independent METABRIC and CGGA cohorts, and evaluated through an explicitly adversarial benchmarking strategy. The study also incorporated orthogonal validation at the proteomic and single-cell levels, assessment of tumour microenvironment and immunotherapy response, drug-sensitivity analysis, and functional enrichment. Importantly, negative and limiting results were retained rather than omitted, including the lack of survival separation among unsupervised PCD subtypes and the lack of immune-checkpoint blockade predictive value.

Several limitations should also be acknowledged. First, the study is entirely computational and based on public datasets; therefore, the conclusions concern statistical behaviour, prognostic performance, and multi-omic consistency rather than direct experimental mechanism. Second, previously published multi-death signatures were not individually benchmarked because uniformly curated model coefficients and gene lists were not consistently available; the main comparators were therefore constituent single-pathway models and random-null signatures. Third, proliferation adjustment relied on the meta-PCNA signature, and alternative proliferation proxies may yield quantitatively different results. Fourth, the immunotherapy cohorts were limited in size, particularly GSE78220, which restricts statistical power for response analyses. Fifth, single-cell validation was performed in one melanoma dataset, and additional single-cell cohorts across other tumour types would strengthen the generality of the observed crosstalk. Finally, although pan-cancer analysis provides broad statistical power, it may obscure lineage-specific biology; for this reason, cancer-specific estimates were also reported where appropriate. In conclusion, the integrated ferroptosis–cuproptosis–disulfidptosis signature captures a real and reproducible biological axis across cancers, but its prognostic value is limited by redundancy with ferroptosis-associated signal and substantial proliferation confounding. The signature stratifies survival but does not predict immune-checkpoint blockade response. These findings argue for a more cautious interpretation of integrated regulated cell-death signatures and support routine incorporation of single-pathway, random-null, and proliferation-adjusted controls in future prognostic biomarker studies.

## 5. Conclusion

The integrated ferroptosis–cuproptosis–disulfidptosis signature captures a reproducible and biologically grounded cell-death crosstalk axis across human cancers. However, its prognostic value is limited by redundancy with the strongest single pathway, substantial proliferation confounding, and lack of predictive performance for immune-checkpoint blockade response. These findings indicate that biological coherence and external reproducibility do not necessarily imply incremental clinical utility.

The crosstalk among ferroptosis, cuproptosis, and disulfidptosis remains an important area for mechanistic investigation and therapeutic exploration [52]. However, its translation into prognostic biomarker models should be evaluated using controls that distinguish genuine added value from reproducible redundancy. The pan-cancer benchmark and openly released code and data provided here offer a reusable framework for assessing future regulated cell-death signatures.

## Declarations

## Author contributions

A.Y.D. and E.Y. conceived and designed the study and developed the analytical framework. A.Y.D. performed data curation, computational analyses, and visualisation, and prepared the original manuscript draft. E.Y. supervised the study, contributed to methodology development and data interpretation, and critically reviewed and edited the manuscript. Both authors read and approved the final version of the manuscript.

## Ethics approval and consent to participate

This study was based exclusively on publicly available, de-identified datasets, including TCGA, METABRIC, CGGA, CPTAC, GEO datasets GSE72056 and GSE78220, IMvigor210, and GDSC2. No new experiments involving human participants, human tissue, animals, or cell lines were performed. Therefore, additional ethics approval and informed consent were not required. Ethical approval and consent procedures were obtained by the original studies contributing these datasets.

## Acknowledgements

The authors gratefully acknowledge the TCGA Research Network, CPTAC, the METABRIC and CGGA consortia, the Genomics of Drug Sensitivity in Cancer project, and the contributors of the GEO and IMvigor210 datasets used in this study.

## Data and code availability

The analysis code, processed result tables, intermediate analysis checkpoints, and large supplementary data files, including Supplementary Data SD1–SD3, are deposited in public repositories. The analysis code will be made available through GitHub, and processed data and supplementary files will be archived on Zenodo.

## Declaration of generative AI and AI-assisted technologies in the writing process

During the preparation of this work the authors used ChatGPT (OpenAI, L.L.C., San Francisco, CA, USA) and DeepL Write (DeepL SE, Cologne, Germany) in order to improve the writing process, including spelling and grammar corrections. After using these tools, the authors reviewed and edited the content as needed and take full responsibility for the content of the publication.

